# The effects of early-life ambient temperature on the gut microbiome of a wild bird? an experimental approach

**DOI:** 10.1101/2024.11.26.625491

**Authors:** Suvi Ruuskanen, Lotta Hollmen, Antoine Stier, Bin-Yan Hsu, Nina Cossin-Sevrin, Coline Marciau, Mikaela Hukkanen, Eero Vesterinen

## Abstract

Vertebrate gut microbiome has significant effects on host development, health, and fitness. Multiple external factors contribute to gut microbiome variation, and the role of ambient temperature has gained increasing attention. Yet, temperature effects are often tested in extremes and in captive systems. We experimentally studied the effect of subtle temperature decreases during post-natal development on gut microbiome diversity and composition in wild pied flycatchers (*Ficedula hypoleuca)*. We also performed partial cross-fostering to study the relative contribution of genetic and rearing environment on microbiome. Nest-box cold treatment did not influence gut microbiome diversity or composition, which may be due to the small temperature change, ontogenetic stage, or other factors, such as diet, causing large variation in the data. Rearing environment explained more of the variation in gut microbiome than genetic background, but the variance explained was relatively small. Future studies need to further address the drivers of the large intraspecific variation in microbiome in natural populations.

## Introduction

The gut microbiome is important for individual health and well known to be involved in host digestion and metabolism (e.g. Chevalier et al. 2015; Cox et al. 2014), immune function (e.g. Knutie et al. 2017; Warne et al. 2019), detoxification processes (e.g. Kohl et al. 2014) and also influences host behavior (e.g. Davidson et al. 2018; Grond et al. 2018). The gut microbiome varies across individuals, populations, and the structure of the microbiome is determined by complex and dynamic interactions between the host, the environment, and microbe-microbe interactions species (McFall-Ngai et al. 2013). In birds, gut microbiome shows very large environmental variation (Bodawatta et al. 2022) which has been associated with e.g. diet (Bodawatta et al. 2021), but importantly, a large proportion of the variation remains unexplained.

Interestingly, several recent vertebrate studies report associations between ambient temperature and the community composition and function of the gut microbiome both in ectotherms but also endotherms (Liu et al. 2023; Sepulveda and Moeller 2020; Zietak et al. 2016, Liukkonen et al. 2024). In birds, data is scarce, but a study on captive chicken exposed to cold stress reported influences on microbiome composition (Yang et al. 2021). Furthermore, microbiome composition in pigeons (*Columba livia*) reared in outdoor aviaries showed seasonal, likely temperature-associated variation (Dietz, et al. 2022) and in passerines, temperature explained small, but significant part of gut microbiome diversity and compositional variation in a continental scale study (Liukkonen et al. 2024). To our knowledge, in wild birds, the effect of postnatal temperature has been studied in only one study: in eastern bluebirds (*Sialis sialis*) and tree swallows *(Tachycineta bicolor*) in Minnesota, moderate warming during the nestling stage tended to lead to lower diversity (Ingala et al. 2021).

Importantly, studies on rodents have revealed the functional consequences of temperature-induced variation in microbiome: when microbiome from chronically cold-exposed mice/voles was transferred to germ-free animals, they showed adaptation to cold, such as increased feed conversion, higher body temperature and metabolic rate, suggesting that the microbiome plays a key role in these adaptive changes (Chevalier et al. 2015; Sepulveda and Moeller 2020; Zhang et al. 2018; Zietak et al. 2016). Furthermore, also in ectotherm tadpoles and insect models (Drosophila), depletion of the gut microbiome reduced the host’s thermal tolerance and survival (Fontaine et al. 2022; Jaramillo and Castañeda 2021). Yet, most previous studies on the topic have been conducted in adult animals, and information on the effects and consequences of early-life temperature on microbiome remain largely unknown. One could argue that in developing animals where temperature-regulation is not fully matured, changes in temperature may play even a larger, and potentially more long-lasting role on their physiology and fitness. Second, most prior studies are from captive species, and using large temperature differences (e.g. reflecting seasonal differences) while it is not understood whether microbiome responds to small-scale temperature variation within a season (Ruuskanen 2024).

To address these knowledge gaps, we experimentally studied the effects of early-life ambient temperature on the development of avian gut microbiome. We manipulated ambient temperature during postnatal period (from 7 to 13 days after hatching) by experimentally decreasing the temperature within nest boxes using a passerine bird, the pied flycatcher (*Ficedula hypoleuca*) nestlings as model system. We measured fecal microbiome diversity and composition before and after the temperature treatment. We predicted that a decrease in temperature will decrease nestling gut microbiome diversity if cooler temperatures act as stressor (similarly as hot temperatures, Ingala et al. 2021; Sepulveda and Moeller 2020). We also expect an overall change in microbial composition between the treatment groups. In addition, we applied a partial cross-fostering design, which allows us to estimate the relative significance of genetic/prenatal and early-postnatal *versus* late post-natal environmental factors in shaping the avian gut microbiome. We predicted that postnatal (foster) nest environment contributes more to microbiome variation than genetic/prenatal factors, as shown in several other passerine studies using cross-fostering designs (Somers et al. 2023; Teyssier et al. 2018; Lucas and Heeb 2005). The effect of local nest environment on microbial communities could be explained by parental effects (incl. vertical transfer of bacteria) and exposure to the same nest microbiome.

## Methods

The study was conducted in a nest-box population (300 boxes) in Ruissalo (60°26’N, 22°10’E), southwestern Finland. The pied flycatcher, is a small, altricial, insectivorous secondary cavity nesting bird, fledging around 16-18 days after hatching (Lundberg and Alatalo 1992). Nests were monitored regularly to record egg-laying and a total of 60 nests were included in this study. Half of the nests were included in an experiment where egg thyroid hormone levels were elevated (similar protocol as in Hsu et al. 2020). This has been accounted for in the statistical models - yet, as hormone manipulation did not influence growth, physiological biomarkers, or survival (Stier et al., unpubl data), it is unlikely to bias the results.

Nest boxes were further monitored to record hatching date. On day 2 after hatching, half of the nestlings (minimum two individuals) were swapped between nests (with nestlings of the same age and weight). Thereafter, each nest contained both biological offspring of the rearing pair, and foster nestlings. This partial cross-fostering design allowed us to separate the relative contribution of genetic background, prenatal and early postnatal environments *versus* later rearing environment (d2 post-hatching onwards). Nestlings were coded with clipping nails for individual identification and weighed on day 2 (mass∼ 0.1g). They were ringed and measured (wing ∼ 0.1mm) on day 7 post-hatching (pre-treatment), and weighed and measured again on day 13 post-hatching (post-treatment).

During days 7-13 post-hatching, half of the nest boxes (N=30) received a cold treatment, and the other half served as controls (N=29). During this period of the nestling stage, mothers no longer regularly brood the nestlings, yet the nestlings are still not fully able to thermoregulate (Lundberg and Alatalo 1992), therefore being more susceptible to environmental temperature variation. Based on pilot experiments, our cold treatment aimed to reduce the nest-box temperature by 2-4 ºC. The experimental protocol was the following: two fabric bags with insulation were stapled to the sides (outer walls) of the nest boxes, and a 600g freezer block (from -20°C) was inserted in each bag. Our pilot trials indicated that the cooling effect lasted ca 8-10h (depending on outside temperature). We thus replaced the blocks every morning to reduce the temperature during the daytime, where the effect of cooling would be strongest. In the control treatment nest boxes, similar fabric bags were attached but contained an unfrozen freezer block. Control nests were similarly visited every day and thus received the same amount of disturbance. To monitor nest box temperature, an iButton (DS1922L-F5, Homechip, UK) temperature logger was attached to the inner side of the back wall, at approximately 5 cm below the ceiling in all treatment and control boxes. Temperature was measured every third minute and daily averages (midnight to midnight) were calculated.

To study the effects of rearing environment and temperature on microbiome, cloacal swab samples were collected both prior to the cold treatment (7 days after hatching) and after (13 days after hatching), using a plastic swab (Copan Floqswabs) moistened with sterile phosphate-buffered saline (PBS) prior for sampling, delicately inserted in the cloaca with a rotational movement for 5 sec. After sampling, the tip of the swab was cut and immediately placed into a sterile 1.5 ml Eppendorf tube, and tubes were stored at -20 °C until DNA extraction. The experiments were conducted under licenses from the Animal Experiment Board of the Administrative Agency of South Finland (ESAVI/5718/2019) and South-Western Finland Centre for Economic Development, Transport and Environment (VARELY 924/2019).

### DNA extraction and library preparation

Due to the large number of samples, we randomly selected samples of one biological and one foster nestling from every nest and analysed repeated samples from the same individuals at the age of 7 and 13 days when available (2 samples at d7 were missing due to chicks being too small for sampling). The final sample sizes were: 118 cool-treated nestling samples (7d N=58 individuals, 13d N=60) and 111 control nestling samples (7d N=56, 13d N=55). Moreover, four negative controls were included, two collected from the field (empty swab in PBS) and two from extraction. The DNA was extracted using the ZymoBIOMICS DNA Microprep Kit (cat nr D4300, Zymo Research, Irvine CA, USA) following manufacturer’s instructions (manual version 2020). Thereafter, the library preparation protocol included two separate PCR stages. First, locus-specific PCR was conducted using universal bacterial primers (Bakt_341F: CCTACGGGNGGCWGCAG, Bakt_805R: GACTACHVGGGTATCTAATCC) that were targeted at V3–V4 region of 16S rRNA gene (Herlemann et al. 2011). The first PCR was performed using 6.25 μL MyTaq RedMix DNA polymerase (Meridian Bio-science; Cincinnati, OH, USA), 0.125 μL (final conc100nM) each primer, 1.5 μL DNA filled with sterile water to the total volume of 12.75 μL. The conditions of the first PCR were as follows: 3 min 95°C, followed by 23 cycles of 30 sec 95°C – 30 sec 55°C – 30sec 72°C, and then 10 min of 72°C. Each sample was amplified in two replicates during the first PCR. For the second, library preparation PCR, 2 μL of diluted 1^st^ PCR product was mixed with 1 μL (final conc 500nM) of each Illumina-compatible adapters (including a unique sample-specific combination of indices), 5 μL of MyTaq RedMix polymerase, and 1 μL of water in a 10 μL reaction volume. Second PCR conditions were: 4 min 95°C, followed by 14 cycles of 20 sec 98°C – 15 sec 60°C – 30sec 72°C, and then 3 min 72°C. The success of the PCR’s was determined in a subsequent electrophoresis. Prior to sequencing, DNA concentrations were quantified using Qubit 2.0 Fluorometer and all samples were pooled in equimolar ratios per replicate (one pool for replicate1, another pool for replicate2) for sequencing. The pools were then purified using Solid Phase Reversible Immobilization (SPRI) beads and then checked and quantified with the Agilent 2100 Bioanalyzer. Sequencing was performed using the Illumina MiSeq Reagent Kit v3 (2×300 bp) by the Finnish Functional Genomics Center (Turku, Finland).

### Data processing

Primers and chimeric sequences were first removed and thereafter processed through DADA2 pipeline (Callahan, McMurdie, and Holmes 2017) in software R (version 4.0.2) to call for amplicon sequence variants (ASVs) following Morrill et al. (2021). Data from the replicates were combined retaining an ASV if it was detected in both replicates. Taxonomy was assigned using RDP version v18 database (downloaded from https://www.drive5.com/usearch/manual/sintax_downloads.html) with USEARCH SINTAX (Edgar 2016). Thereafter we conducted multiple filtering steps: 1) any ASV that could not be assigned to any taxa, were discarded, 2) any ASV with fewer than 2 reads were discarded, and 3) ASVs were removed from a sample, if negative controls had higher abundance of particular ASV than samples. For the analysis of alpha diversity, the data was rarefied to 1,000 sequences per sample, resulting in the loss of 39 samples with fewer than 1,000 sequences. Consequently, the sample size in each treatment group was: 98 cool-treatment samples (7d N=51, 13d N=47) and 92 control samples (7d N=48, 13d N=44).

### Statistical analyses

The statistical analyses were conducted with R software (version 4.0.2). Daily averages of temperatures from d7 to d13 in each nest box were calculated, and temperature differences between cool-treatment group and control group were analyzed using a linear mixed model (package *lme4* Bates et al. 2015) with group (treatment or control) as the fixed factor, and nest box identity as random factor.

Microbiome alphadiversity (using function *alpha* in package *microbiome*, Lahti and Shetty 2012), was characterized using observed richness/observed ASVs, Shannon index (H’), and Pielou evenness index (J). Linear mixed models were used where cold treatment, age (7 or 13 days), cross-fostering (cross-fostered or not, to account for the act of moving the chicks between nests), the number of 2-day-old chicks in rearing nest (to control for effects of brood size), and cold×age interaction were included as fixed effects. The effect of TH treatment was initially tested but dropped from the models as it was non-significant. This modelling strategy was chosen as we wanted to control for initial differences, and to explore potential age effects. As the same nestling was sampled at day 7 and 13, nestling identity (ring number) was included as a random effect. Furthermore, the identity of the original nest box (where a chick hatched, i.e nest of origin) and the identity of the rearing nest box (where a chick was reared, i.e. nest of rearing) were included as random effects, to control for non-independence of nestmates, and to study the relative importance of genetic and rearing environment. Note that the estimate for individual identity appeared as zero, suggesting that the effect is very small or that there is singular fit due to missing repeated values. However, due to the sampling design the random effect should be retained in the model. To study microbiome composition (i.e. betadiversity), we conducted a non-metric distance scaling (NMDS) ordination based on Bray-Curtis dissimilarity indices. We used the permutational analysis of variance (PERMANOVA) to estimate the statistical significance of differences between the treatment groups using the *vegan* package and the function *adonis*. PERMANOVA was conducted on data from 13-day individuals only, ie. post-treatment data, with treatment as the only fixed factor as interaction term and repeated sampling could not be included in the model.

## Results

The cold treatment reduced the nest box temperature approximately by 1.3 ºC, resulting in significant difference in nest box temperatures between control and treatment groups (F_1,57_= 7.640, p=0.008, Suppl Fig 1). The average temperature (marginal means ±SE) in a treatment group was 19.3 ±0.31 ºC and in a control group 20.6 ±0.32 ºC during the 7-day temperature manipulation.

In the pied flycatcher fecal microbiome data, overall, the four most abundant bacterial phyla were Proteobacteria (average±SE: 65.6±2.2%), Firmicutes (18.8±1.9%), Actinobacteria (5.8±0.7%), and Chlamydiae (5.8±1.2%, see also Fig 2a). There was no impact of the cold treatment on three alpha diversity indices (Fig 1, Table 1, age*treatment interaction). Alpha diversity did not change with age (Shannon 7d: marginal mean±SE 1.84±0.11, 13d: 1.82±0.10; observed richness 7d: 25.68±3.06, 13d: 23.34±2.87; Pielou evenness 7d: 0.66±0.03, 13d: 0.67±0.02), brood size or between moved and non-moved nestlings (Table 1). The nest of rearing explained some of the variation in Shannon index (17%) and observed richness (5%) while nest of origin contributed very little or none to the variation (Table 1). Cold treatment did not lead to differences in microbial compositions between treatment and control group (d13 data only included: PERMANOVA F=0.832_1,89_, p=0.708, Fig. 2a,b).

**Table 1.** Linear mixed model (LMM) on the effects of cold treatment and covariates on different alpha diversity indices (Shannon, observed richness, and Pielou evenness). The subscripts of F-values represent the numerator and denominator degrees of freedom. Note that as both pre and post-treatment microbiome measurements were included, age*cold treatment interaction tests for the main question, differential effect of cold treatment on diversity.

|  | Shannon index (H') |  | Observed |  | Pielou evenness (J) |  |
| --- | --- | --- | --- | --- | --- | --- |
| Fixed effects | F-value | P-value | F-value | P-value | F-value | P-value |
| Cold treatment | 0.150<sub>1,55</sub> | 0.700 | 0.593<sub>1,51</sub> | 0.445 | 1.953<sub>1,47</sub> | 0.169 |
| Cross-fostering | 0.666<sub>1,172</sub> | 0.416 | 0.001e-01<sub>1,179</sub> | 0.993 | 0.535<sub>1,173</sub> | 0.466 |
| Age | 0.019<sub>1,141</sub> | 0.891 | 0.290<sub>1,142</sub> | 0.591 | 0.071<sub>1,137</sub> | 0.790 |
| Brood size (d2) | 1.313<sub>1,57</sub> | 0.257 | 2.842<sub>1,54</sub> | 0.098 | 0.036<sub>1,49</sub> | 0.851 |
| Cold*Age | 0.003<sub>1,141</sub> | 0.955 | 0.097<sub>1,142</sub> | 0.756 | 0.092<sub>1,137</sub> | 0.762 |
| Random effects | Variance |  | Variance |  | Variance |  |
| Individual | 0.000 |  | 0.000 |  | 0.000 |  |
| Nest of origin | 0.000 |  | 1.375e-07 |  | 3.274e-11 |  |
| Nest of rearing | 0.193 |  | 44.08 |  | 0.003 |  |
| Residual | 0.919 |  | 799.50 |  | 0.054 |  |

**Figure 1.**
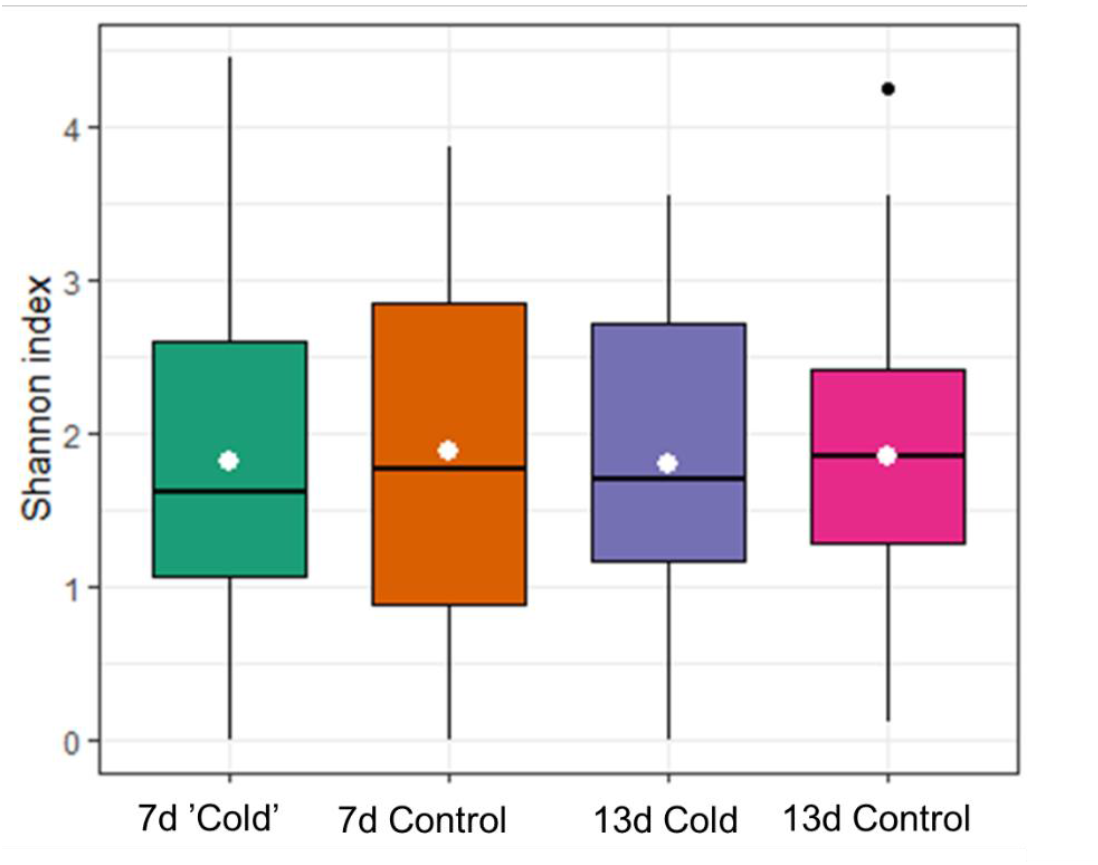
Shannon indices of four different age*treatment groups. 7d nestlings have been sampled prior to cold treatment and 13d nestlings after cold treatment. Median is represented by the black crossbar, the mean by the white dot and 25 % and 75 % quartiles by the lower and upper box. Whiskers show the range between minimum and maximum value, excluding one outliers represented in black dot. Sample sizes: 7d cold N=28 nests/51 nestlings, 7d control N=29/48, 13d cold N=30/47, 13d control N=26/44.

**Figure 2.**
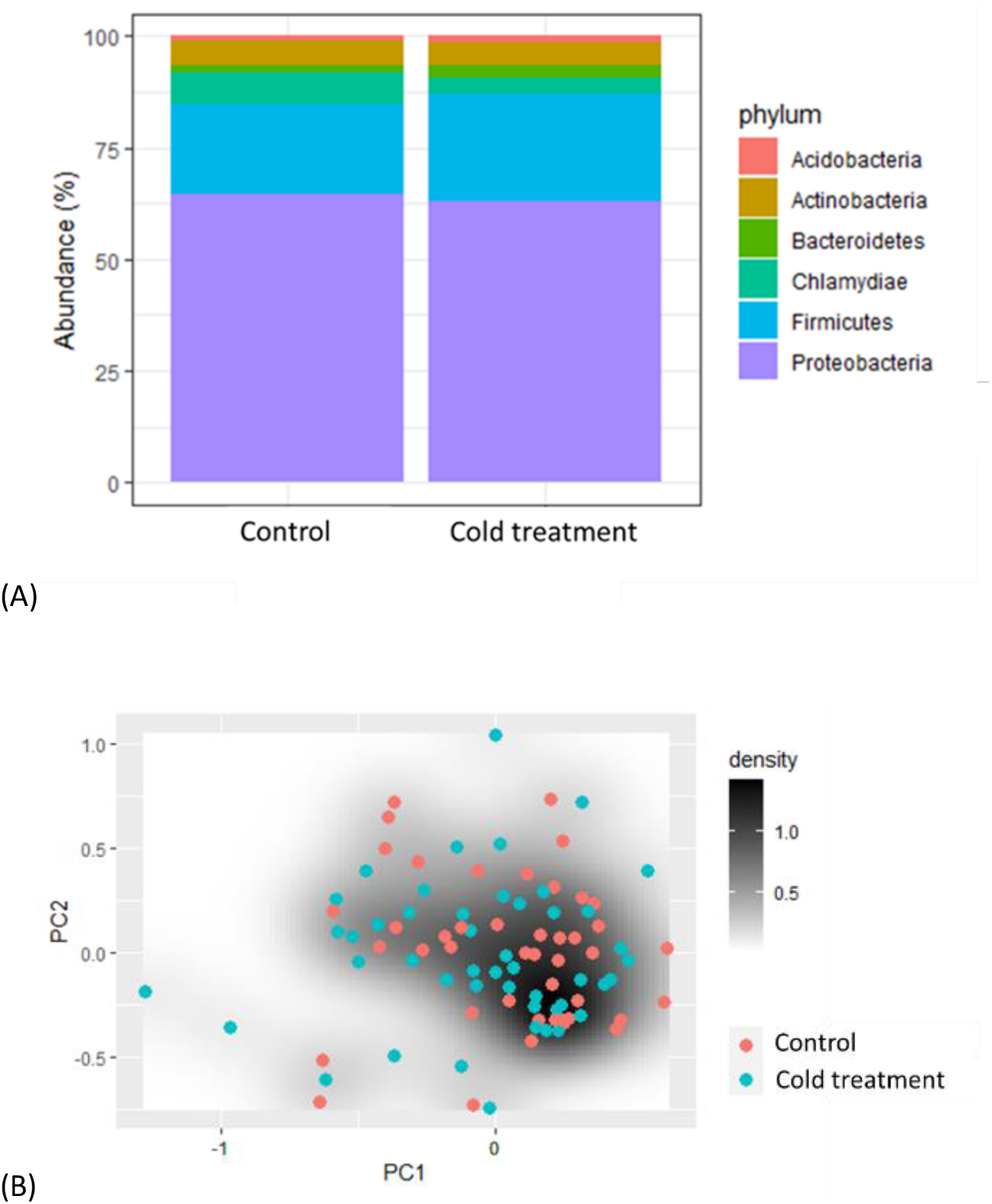
(A). The average relative abundances of main bacterial phyla in control and cold treatment groups of 13 day-old pied flycatcher nestlings, i.e. post-treatment samples. Sample sizes: cold treatment N=30 nests/47 nestlings, control N=26 nests/44 nestlings. (B) Principal coordinate analysis (PCoA) based on Bray-Curtis dissimilarity between cold treatment (blue) and control (red) groups. Sample sizes: cold treatment N=30 nests/47 nestlings, control N=26 nests/44 nestlings. The analysis included only 13 day-old pied flycatcher nestlings, ie. post-treatment samples.

## Discussion

Our results suggest that short-term, mild temperature decrease does not influence cloacal microbiome diversity or composition in a developing altricial bird. The lack of effect of the experimental cooling on gut microbiome diversity and composition may be explained by several factors: (1) The experimental cooling led to an average temperature decrease of 1.3°C, which was a relatively small but still a statistically significant difference. Previous thermal experiments that have reported significant changes in microbiome diversity/composition have used considerably larger temperature differences (5-10°C, Sepulveda and Moeller 2020). The potential mechanisms underlying ambient-temperature effects in these previously published studies are not fully understood: changes in microbiome may stem from other physiological changes of the host (e.g. gut physiology or stress physiology), while especially in ectotherms they may reflect direct changes in body temperature and therefore the actual temperature microbes are exposed to (Sepulveda and Moeller 2020). These physiological responses were not measured in our study, and it would be important to take that into account in future studies to try to disentangle the mechanisms via which microbiome may or may not be affected by ambient temperature, especially in endotherms. Yet, our results suggest that the small changes in ambient thermal environment as in this experiment may not lead to major effects on physiology that would cause changes in offspring’s gut microbiome. (2) The majority of the prior data originates from captive organisms where the microbiome is heavily influenced by captivity (also in birds, e.g. Kelly et al. 2022). In wild populations, other environmental variation such as variation in diet may have overridden the subtle effects of temperature manipulation. However, wild studies are needed to understand the complexity of gut microbiome variation and the associated phenotypes in natural environments. (3) The effect of subtle temperature changes may depend on the ontogenetic stage, which is associated with the thermoregulatory capacity and the stability of the microbiome. Here, ambient temperature was manipulated when thermoregulatory capacity was developing (day 7 onwards; Rodriguez, Diez-Mendez, and Barba 2016). Therefore we cannot conclude whether temperature changes influence gut microbiome in very young nestlings (less than 7d old) totally incapable of thermoregulating, and in which the microbiome is changing more rapidly (Teyssier et al. 2018; Somers et al. 2023). Nevertheless, pied flycatcher females brood their offspring until the age of 7 days (Sanz and Moreno 1995), and therefore an experiment before the age of 7 would have been confounded by the heat provided by the brooding female. Finally, to our knowledge the only previous study on the effects of temperature on developing altricial birds in wild populations similarly showed that temperature increase (+5°C) had only small effects on gut microbiome (Ingala et al. 2021).

Cross-fostering in our experimental setup enabled the investigation of the roles of genetic background and prenatal environment compared to the later rearing environment in shaping nestlings’ gut microbiome. Genetic background could influence gut microbiome for example via influencing host gut physiology or immune function (e.g. Blekhman et al. 2015). Parallel to our predictions, the rearing environment explained more of the variation than genetic background, yet, the effect of rearing nest on gut microbiome diversity was relatively small (ca 5-17%). Previous microbiome studies on passerines have indeed shown quite clearly that foster siblings living in the same nest share more similar gut microbiome than biological siblings inhabiting different nests (Somers et al. 2023; Teyssier et al. 2018; Lucas and Heeb 2005). The effect of the nest environment on microbial communities has been explained by parental effects (including horizonal transfer), transfer of bacteria between nestlings and from the nest material (Lucas & Heeb 2005; Teyssier et al. 2018). Importantly, some of the differences between the studies may arise due to methodological differences, as both fecal and cloacal swabs have been used in the studies of the gut microbiome.

All in all, we found large individual variation in microbiome diversity and composition, which was not explained by subtle changes in early-life temperature. The nest rearing environment (but not genetic background) explained a small proportion of the variation, yet there is a large proportion of unexplained intraspecific variation in gut microbiome in birds in our study and bird studies in general. The drivers of this (dynamic) variation should be characterized, and their phenotypic associations and fitness consequences should be explored in future studies.

## Acknowledgements

We thank all field assistants, Axelle Delauany and Melanie Cromebque, for their valuable help in data collection. We also thank Meri Lindqvist for excellent help in the laboratory analyses and troubleshooting.

## Author contributions

Conceptualization: S.R., E.V., L.H. Methodology: S.R., A.S., B.Y.H. ;

Formal analysis: L.H., E.V., S.R. Investigation: S.R., L.H., A.S., B.Y.H, N.C.S, C.M. and M.H., Resources: S.R., E.V.; Data curation: E.V., L.H.; Writing - original draft:

S.R.; Writing - review & editing: S.R., L.H., A.S., B.Y.H, N.C.S, C.M. and M.H.;

Supervision: S.R., E.V.; Project administration: S.R.; Funding acquisition: S.R., A.S., E.V.

## Data availability

Data will available in the Dryad Digital repository LINK

## Supplementary material

**Fig 1.**
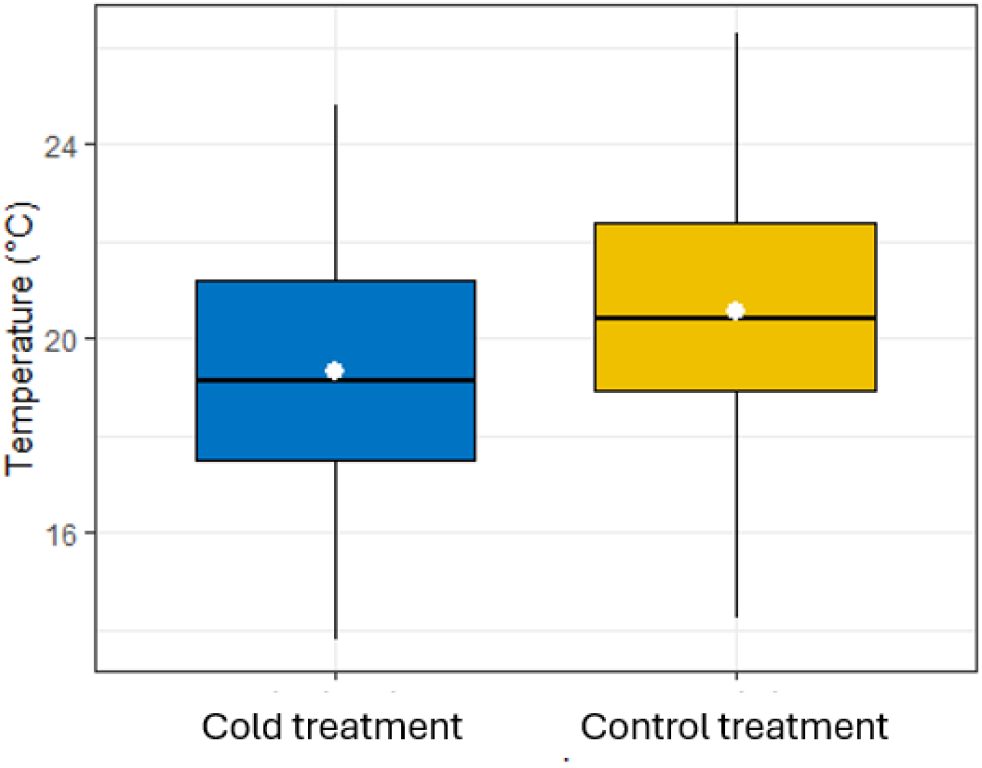
Average temperature inside nest boxes overaged over the cold treatment period (d7-d13 post-hatching). Sample sizes Ncold = 30, Ncontrol = 29.

## References

Bates, D., M. Machler, B. M. Bolker, and S. C. Walker. 2015. Fitting Linear Mixed-Effects Models Using Lme4. Journal of Statistical Software 67(1): 1–48. doi:10.18637/jss.v067.i01.

Blekhman, R., Goodrich, J.K., Huang, K. et al. 2015. Host genetic variation impacts microbiome composition across human body sites. Genome Biologyl 16:191. doi: 10.1186/s13059-015-0759-

Bodawatta, K.H., Freiberga, I., Puzejova, K., Sam, K., Poulsen, M., and Jønsson, K.A. 2021. Flexibility and Resilience of Great Tit (Parus Major) Gut Microbiomes to Changing Diets. Animal Microbiome 3(1): 20. doi:10.1186/s42523-021-00076-6.

Bodawatta, K H., Hird, S.M., Grond, K., Poulsen, M. and Jønsson, K.A. 2022. Avian Gut Microbiomes Taking Flight. Trends in Microbiology 30(3): 268–80. doi:10.1016/j.tim.2021.07.003.

Callahan, B.J., McMurdie, P.J., and Holmes, S.P. 2017. Exact Sequence Variants Should Replace Operational Taxonomic Units in Marker-Gene Data Analysis. The ISME Journal 11(12): 2639–43. doi:10.1038/ismej.2017.119.

Chevalier, C., O. Stojanovic, D. J. Colin, N. Suarez-Zamorano, V. Tarallo, C. Veyrat-Durebex, D. Rigo, et al. 2015. Gut Microbiota Orchestrates Energy Homeostasis during Cold. Cell 163(6): 1360–74. doi:10.1016/j.cell.2015.11.004.

Cox, L.M. et al. 2014, Altering the intestinal microbiota during a critical developmental window has lasting metabolic consequences. Cell 158:705–721

Davidson, G.A., Cooke, A.C., Johnson, C.N, Quinn, J.L. 2018. The gut microbiome as a driver of individual variation in cognition and functional behaviour. Philos. Trans. R. Soc. B. Biol. Sci. 373: 20170286–20170286.

Dietz, M.W., Matson, K.D., Versteegh, M.A., van der Velde, M., Parementier, H.K., Arts, J.A.J., Salles, J.F., Tieleman, I.B. 2022. Gut Microbiota of Homing Pigeons Shows Summer-Winter Variation under Constant Diet Indicating a Substantial Effect of Temperature. Animal Microbiome 64

Edgar, R.C. 2016. SINTAX: A Simple Non-Bayesian Taxonomy Classifier for 16S and ITS Sequences. : 074161. doi:10.1101/074161.

Fontaine, S.S., Mineo, P.M and Kohl, K.D. 2022. Experimental Manipulation of Microbiota Reduces Host Thermal Tolerance and Fitness under Heat Stress in a Vertebrate Ectotherm. Nature Ecology & Evolution 6(4): 405–17. doi:10.1038/s41559-022-01686-2.

Herlemann, D.P.R, Labrenz, M., Jürgens, K., Bertilsson, S., Waniek, J.J. and Andersson, A.F. 2011. Transitions in Bacterial Communities along the 2000 Km Salinity Gradient of the Baltic Sea. The ISME Journal 5(10): 1571–79. doi:10.1038/ismej.2011.41.

Hsu, B. Y., T. Sarraude, N. Cossin-Sevrin, M. Crombecque, A. Stier, and S. Ruuskanen. 2020. Testing for Context-Dependent Effects of Prenatal Thyroid Hormones on Offspring Survival and Physiology: An Experimental Temperature Manipulation. Scientific Reports 10(1). doi:10.1038/s41598-020-71511-y.

Ingala, M.R., Albert, L., Addesso, A., Watkins, M.J. and Knutie, S.A. 2021a. Differential Effects of Elevated Nest Temperature and Parasitism on the Gut Microbiota of Wild Avian Hosts. Animal Microbiome 3(1): 67. doi:10.1186/s42523-021-00130-3.

Jaramillo, A. and Castañeda, LW. 2021. Gut Microbiota of Drosophila Subobscura Contributes to Its Heat Tolerance and Is Sensitive to Transient Thermal Stress. Frontiers in Microbiology 12: 654108. doi:10.3389/fmicb.2021.654108.

Kelly, T.R, Vinson, A.E., King G.M. and Lattin, C.R. 2022. No Guts About It: Captivity, But Not Neophobia Phenotype, Influences the Cloacal Microbiome of House Sparrows (Passer Domesticus). Integrative Organismal Biology 4(1): obac010. doi:10.1093/iob/obac010.

Liu, S., Xiao Y., Wang, X., Guo, D., Wang, Y. and Wang, Y. 2023. Effects of Microhabitat Temperature Variations on the Gut Microbiotas of Free-Living Hibernating Animals. Microbiology Spectrum 11(4): e00433–23. doi:10.1128/spectrum.00433-23.

Liukkonen, M., Muriel, J., Martinez-Padilla, J., Nord, A., Pakanen, V-M., Rosivall, B., Tilgar, V., van Oers, K., Grond, K., Ruuskanen, S. 2024. Seasonal and environmental factors contribute to the variation in the gut microbiome: A large-scale study of a small bird. Journal of Animal Ecology. DOI: 10.1111/1365-2656.14153

Lucas, F.S. and Heeb, P. 2005. Environmental Factors Shape Cloacal Bacterial Assemblages in Great Tit Parus Major and Blue Tit P. Caeruleus Nestlings. Journal of Avian Biology 36(6): 510–16. doi:10.1111/j.0908-8857.2005.03479.x.

Lundberg, A., and R. Alatalo. 1992. The Pied Flycatcher.

Poyser Kohl, K.D., Weiss, R.B., Cox, J., Dale, C., Dearing, D.M. 2014. Gut microbes of mammalian herbivores facilite intake of plant toxins. Ecology Letters 17: 1238–1246. doi: 10.1111/ele.1232

Knutie, S.A., Wilkinson, C.L., Kohl, K.D., Rohr, J.R. 2017. Early-life disruption of amphibian microbiota decreases later-life resistance to parasites. Nature Communications. 8:86.

Warne, R.W., Kirschman, L.J., Zeglin, L. 2019. Manipulation of gut microbiota during critical developmental windows affects host physiological performance and disease susceptibility across ontogeny. Journal of Animal Ecology 88, 845–856.

McFall-Ngai, M., M. G. Hadfield, T. C. G. Bosch, H. V. Carey, T. Domazet-Loso, A. E. Douglas, N. Dubilier, et al. 2013. Animals in a Bacterial World, a New Imperative for the Life Sciences. Proceedings of the National Academy of Sciences of the United States of America 110(9): 3229–36. doi:10.1073/pnas.1218525110.

Morrill, A., Kaunisto, K.M., Mlynarek, J.J., Sippola, E., Vesterinen, E.J. and Forbes, M.R. 2021. Metabarcoding Prey DNA from Fecal Samples of Adult Dragonflies Shows No Predicted Sex Differences, and Substantial Inter-Individual Variation, in Diets. PeerJ 9: e12634. doi:10.7717/peerj.12634.

Rodriguez, S., D. Diez-Mendez, and E. Barba. 2016. Negative Effects of High Temperatures during Development on Immediate Post-Fledging Survival in Great Tits Parus Major. Acta Ornithologica 51(2): 235–44. doi:10.3161/00016454ao2016.51.2.009.

Ruuskanen, S. 2024. Early-Life Environmental Effects on Birds: Epigenetics and Microbiome as Mechanisms Underlying Long-Lasting Phenotypic Changes. Journal of Experimental Biology 227(Suppl_1): jeb246024. doi:10.1242/jeb.246024.

Sanz, J. J., and J. Moreno. 1995. EXPERIMENTALLY-INDUCED CLUTCH SIZE ENLARGEMENTS AFFECT REPRODUCTIVE SUCCESS IN THE PIED-FLYCATCHER. Oecologia 103(3): 358–64. doi:10.1007/bf00328625.

Sepulveda, J., and A. H. Moeller. 2020. The Effects of Temperature on Animal Gut Microbiomes. Frontiers in Microbiology 11. doi:10.3389/fmicb.2020.00384.

Somers, S.E., Davidson, G.L., Johnson, C.N., Reichert, M.S., Crane, J.M.S., Ross, J.P., Stanton C. and Quinn, J.L. 2023. Individual Variation in the Avian Gut Microbiota: The Influence of Host State and Environmental Heterogeneity. Molecular Ecology n/a(n/a). doi:10.1111/mec.16919.

Teyssier, A., L. Lens, E. Matthysen, and J. White. 2018. Dynamics of Gut Microbiota Diversity During the Early Development of an Avian Host: Evidence From a Cross-Foster Experiment. Frontiers in Microbiology 9: 12. doi:10.3389/fmicb.2018.01524.

Yang, Y., Gao, H., Li, X., Cao, Z., Li, M., Liu, J., Qiao, Y. et al. 2021. Correlation Analysis of Muscle Amino Acid Deposition and Gut Microbiota Profile of Broilers Reared at Different Ambient Temperatures. Animal Bioscience 34(1): 93–101. doi:10.5713/ajas.20.0314.

Zhang, X. Y., G. Sukhchuluun, T. B. Bo, Q. S. Chi, J. J. Yang, B. Chen, L. Zhang, and D. H. Wang. 2018. Huddling Remodels Gut Microbiota to Reduce Energy Requirements in a Small Mammal Species during Cold Exposure. Microbiome 6. doi:10.1186/s40168-018-0473-9.

Zietak, M., P. Kovatcheva-Datchary, L. H. Markiewicz, M. Stahlman, L. P. Kozak, and F. Backhed. 2016. Altered Microbiota Contributes to Reduced Diet-Induced Obesity upon Cold Exposure. Cell Metabolism 23(6): 1216–23. doi:10.1016/j.cmet.2016.05.001.

